# ColBuilder: Flexible structure generation of crosslinked collagen fibrils

**DOI:** 10.1101/2024.12.10.627782

**Authors:** Debora Monego, Matthias Brosz, Johanna Buck, Vsevolod Viliuga, Jaewoon Jung, Torsten Stuehn, Matthias Schmies, Yuji Sugita, Frauke Gräter

## Abstract

Collagen fibrils are fundamental building blocks of connective tissues, yet generating accurate molecular models of their structure remains challenging due to their hierarchical organization and complex crosslinking patterns. ColBuilder has been developed to automate the generation of atomistic models of crosslinked collagen fibrils and facilitate the setup of molecular simulations. The tool integrates homology modeling, higher-order structure generation and optimization to build complete fibril structures with precise control over sequence composition, crosslinking patterns, and dimensions. Users can explore different collagen sequences, manipulate crosslink chemistry through mixed ratios and densities, and generate fibrils of varying diameter and length. All-atom molecular dynamics simulations of 335 nm-long fibrils validate the generated structures, showing excellent agreement with experimental measurements of D-band periodicity and force-extension behavior. ColBuilder is available both as an open-source command-line application and through a web interface at colbuilder.mpip-mainz.mpg.de.

## Introduction

Collagen fibrils are integral components of the extracellular matrix, playing a crucial role in the structural integrity and mechanical properties of various tissues. Understanding the structural and functional dynamics of collagen fibrils is essential for advancing tissue engineering strategies and elucidating mechanisms of collagen-related diseases. However, this understanding remains challenging due to the hierarchical complexity and intricate assembly processes of collagen fibrils [1, 2]. While experimental techniques such as X-ray crystallography [3] and nuclear magnetic resonance (NMR) [4, 5] have provided valuable atomic-level information, they often struggle to resolve the full complexity of collagen structures, particularly in capturing heterogeneity and flexibility.

Computational modeling and molecular dynamics (MD) simulations have emerged as powerful complementary tools to address these limitations. These approaches enable detailed exploration of collagen fibril conformations [6,7], dynamics [8, 9], and interactions [10, 11]. However, preparing accurate initial models for such simulations can be labor-intensive and error-prone, often requiring the integration of multiple tools, data sources, and specialized expertise. While dedicated tools exist for generating models of specific biological systems, such as CHARMM-GUI and Packmol for membranes [12, 13] and nanodiscs [14], there has been a notable absence of analogous tools for filamentous proteins like collagen.

To address this gap, some of us previously developed ColBuilder, a database offering pre-compiled atomistic models of the D-band region of the collagen I fibril [15]. We now introduce the tool ColBuilder, which significantly expands upon its predecessor’s capabilities by enabling end-to-end generation of complex collagen fibrils. ColBuilder implements a template-based modeling strategy, using experimentally derived collagen structures as a foundation for generating diverse fibril models. Key advancements include: (1) homology modeling capabilities for collagens from different species and compositions, (2) advanced crosslinking manipulation, allowing users to mix different crosslinks at varying ratios or control crosslinks density, and (3) full control over the final fibril’s diameter and length. These features directly address the limitations of current computational approaches by providing a streamlined, accurate, and flexible tool for collagen fibril modeling.

ColBuilder is provided as both an open-source command line application built with Python (github.com/graeter-group/colbuilder) and an accompanying web interface (colbuilder.mpip-mainz.mpg.de). To facilitate the use of our models in MD simulations, we also provide a topology generation option and Amber force field parameters for different lysine-derived crosslink types. We have conducted a comprehensive validation study using computational and experimental benchmarks to assess the accuracy and reliability of ColBuilder’s output. The versatility of ColBuilder extends its value beyond MD simulations, with potential applications in Cryo-Electron Microscopy (Cryo-EM) model fitting and as templates for protein design. Finally, we hope that the approach we used in ColBuilder will be transferable to model other filamentous protein materials, such as fibronectin and intermediate filaments.

## ColBuilder: Methodology and Results

Figure 1 illustrates the workflow of ColBuilder for collagen fibril generation. The primary input is a 3D structure of a single collagen triple helix in PDB format, including unit cell symmetry information. ColBuilder offers two main pathways: (1) direct fibril generation, where users specify desired dimensions and crosslinking parameters, and (2) sequence modification and modeling, where users can input custom crosslinking specifications or alter the template’s amino acid sequence. In both cases, the resulting higher-order structure is optimized within a Bravais lattice, with appropriate crosslinking of triple helices.

**Figure 1.**
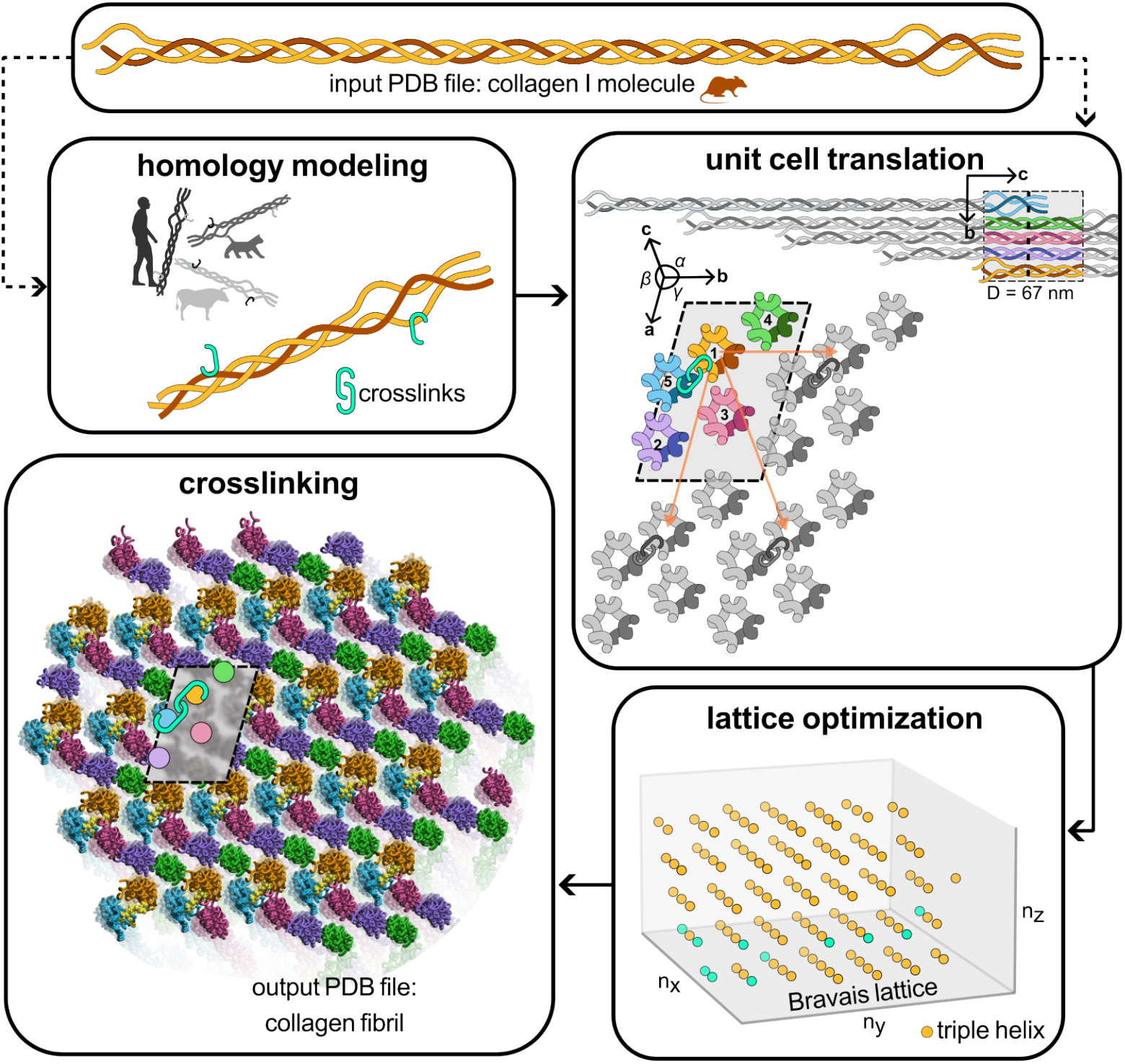
Workflow diagram of ColBuilder, showing the main steps including choice of input structure, generation of copies and their translation to form the higher-order structure, lattice optimization in Bravais lattice and crosslinking to form the final optimized collagen fibril model.

### Homology Modeling and Initial Structure Generation

ColBuilder accommodates two distinct input pathways, each initiating a different processing workflow. Users can either use a previously optimized model of collagen I in PDB format and containing collagen fibril’s unit cell definition, or input a custom collagen amino acid sequence in the FASTA format. In the case of direct structure input, ColBuilder utilizes the provided structure without the need for homology modeling. However, when an amino acid sequence is provided, the software employs a homology modeling process to generate the initial structure.

For amino acid sequence inputs, our homology modeling process uses a template structure of rat collagen type I, derived from the model used in the previous version of ColBuilder [15]. The process begins with a two-step alignment procedure implemented using Biopython [16] to preserve the structural integrity of the collagen triple helix. First, we align the three chains within the template collagen molecule, ensuring the conservation of the characteristic Gly-X-Y repetitive pattern; this aligned triple helix template is then aligned with the target sequence using MUSCLE v3.8.31 [17]. This method allows for the introduction of species-specific variations or desired mutations while maintaining the core collagen structure.

MODELLER performs the homology modeling using a refinement protocol that prioritizes backbone stability. This approach substitutes amino acids without significantly altering the overall backbone conformation, which is critical for maintaining the triple helical structure characteristic of collagen. We validated this procedure for 18 different species with a range of sequence identities (≈ 70 − 98%) to the rat collagen I template. Structural consistency was assessed by per-residue RMSD calculations and preservation of key collagen motifs, particularly the Gly-X-Y pattern. For all sequences tested, the per-residue RMSD was consistently below 2 Å and the average axial rise per triplet (i.e., the distance between consecutive Gly-C^*α*^ in the triple helix) is in agreement with the expected value of approximately 8.6 Å for collagen triple helices [18] (see Supplementary Information, Figure S1 and Table S1), demonstrating the robustness of our method.

ColBuilder’s homology modeling feature greatly enhances its versatility compared to its predecessor, enabling users to generate and analyze collagen structures from different species or with specific mutations. While our approach produces accurate molecular-level structures, we advise users to carefully examine models generated from sequences that deviate significantly from our template (rat collagen, type I). Additionally, it is important to note that fibril-level organization may vary between species, as experimental structural data at this scale is primarily derived from native rat type I collagen [3].

### Higher-Order Structure Generation

Building upon the initial collagen template, ColBuilder generates higher-order structures using UCSF Chimera v1.15’s crystal contacts command [19]. The process begins with a collagen molecule coordinate file in PDB format, defining atoms *A* = *a*_1_, …, *a*_*N*_ at positions ***Q*** = (***q***_1_, …, ***q***_*N*_), along with a user-specified contact distance. The algorithm extracts crystallographic information from the input file, including lattice parameters (*a, b, c, α, β, γ*), space group (*G*_*SP*_), and crystal orientation matrix ***C***. This information defines the unit cell and guides the generation of the higher-order fibril structure. ColBuilder then generates multiple symmetry copies of the unit cell using Euclidean transformation matrices ***T*** = (***R, t***), where ***R*** and ***t*** represent rotation and translation, respectively. Specifically, each atom in the new set *A*^*′*^ undergoes a transformation according to the equation:

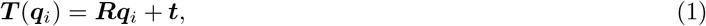

where *i* = 1, …, *N*. The new symmetry copies have positions 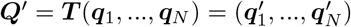 [20].

The contact distance parameter (*dc*) in UCSF Chimera’s crystal contacts tool determines the cutoff distance for including symmetry-related copies of the unit cell. As this distance increases, more symmetry operations are applied, incorporating unit cells that are further from the origin in the crystal lattice. This results in the generation of larger fibril structures. The exact relationship between contact distance and fibril diameter is complex, depending on the unit cell parameters, space group, and specific symmetry operations of the collagen crystal structure. Figure S2 in the Supplementary Information shows the empirical relationship between the contact distance parameter and the resulting fibril radius generated by ColBuilder, demonstrating how users can control the size of the generated microfibril structure by adjusting this parameter. We note that for contact distances *dc* ≤ 15, only up to a couple of complete unit cells are included, and increasing *dc* has no effect in the overall fibril diameter.

A critical step in the higher-order structure generation process is the identification and removal of steric clashes between atoms from different symmetry copies. Our algorithm ensures physically realistic structures by considering the van der Waals radii (*r*_*w,i*_, *r*_*w,j*_) of interacting atoms. It generates a coordinate file of the higher-order structure and transformation matrices for each symmetry copy, enabling subsequent refinement on a discrete Bravais lattice. The resulting system is an ensemble of models *µ* = (*µ*_1_, …, *µ*_*M*_), where each model *µ*_*m*_ contains information about its crosslinks 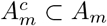 (if present), Euclidean transformation ***T***_*m*_, and Bravais lattice point *p*_*m*_. Any crosslink present in the model is characterized by type (divalent or trivalent; for details, see next section), residue ID, residue name, chain ID, and atomic positions 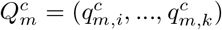 in Cartesian coordinates.

### Structure Optimization

To efficiently represent the quasi-crystalline nature of collagen fibrils, we map higher-order structures from Cartesian space (ℝ^3^) to a discrete lattice ((*n*_*x*_, *n*_*y*_, *n*_*z*_) ∈ ℤ^3^) using the crystal orientation matrix ***C*** (Equation 2). Transformations between spaces are performed using ***p*** = ***C***^−1^***t***, where ***t*** represents the Cartesian translation vector and ***p*** the corresponding lattice point.

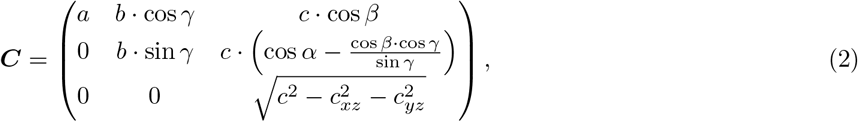

Initial structural analysis of the microfibril revealed a centrosymmetrical pattern with an inversion center at the lattice origin, consistent with the triclinic unit cell of collagen. However, we observed a 35% higher point density within 10 nm of the origin compared to regions beyond 50 nm, indicating denser packing of collagen triple helices near the center. This is a consequence of the finite size of the fibril, resulting in less triple helices at the boundaries of the fibril. Connectivity analysis showed that 73% of molecules were crosslinked at one end, 18% at both ends, and 9% remained unconnected, highlighting the need for structural optimization.

To address these issues, we developed a layer-by-layer optimization approach on the Bravais lattice. The process begins by defining the solution space using *δ* = (*δ*_*x*_, *δ*_*y*_, *δ*_*z*_), which restricts possible solutions to a finite set (Figure 2A, light yellow box). The *δ*_*z*_ parameter selects the number of *n*_*x*_, *n*_*y*_-layers for optimization. For instance, setting *δ*_*z*_ = 2 defines the solution space as the two upper (*n*_*z*,min_ + *δ*_*z*_, *n*_*z*,min_ + *δ*_*z*_ − 1) and two lower (*n*_*z*,max_ − *δ*_*z*_, *n*_*z*, max_ − *δ*_*z*_ + 1) layers from ***l***_***z***_ for optimization. For each layer *l*_*z*_ ∈ ***l***_***z***_, we define a set of points ***P***_***z***_ = (***p***_***i***,***x***_, ***p***_***i***,***y***_, *l*_*z*_) with *i* = 1, …, *N*_*z*_. We then determine the extreme values in the *n*_*x*_, *n*_*y*_ dimensions to create vectors ***l***_***x***_ and ***l***_***y***_, which span a rectangular plane ***P***_***z***,rec_ = (***l***_***x***_, ***l***_***y***_, *l*_*z*_) (Figure 2A, light purple squares). The optimization space is defined as ***P***_***z***,opt_ = ***P***_***z***,rec_ \ ***P***_***z***_.

**Figure 2.**
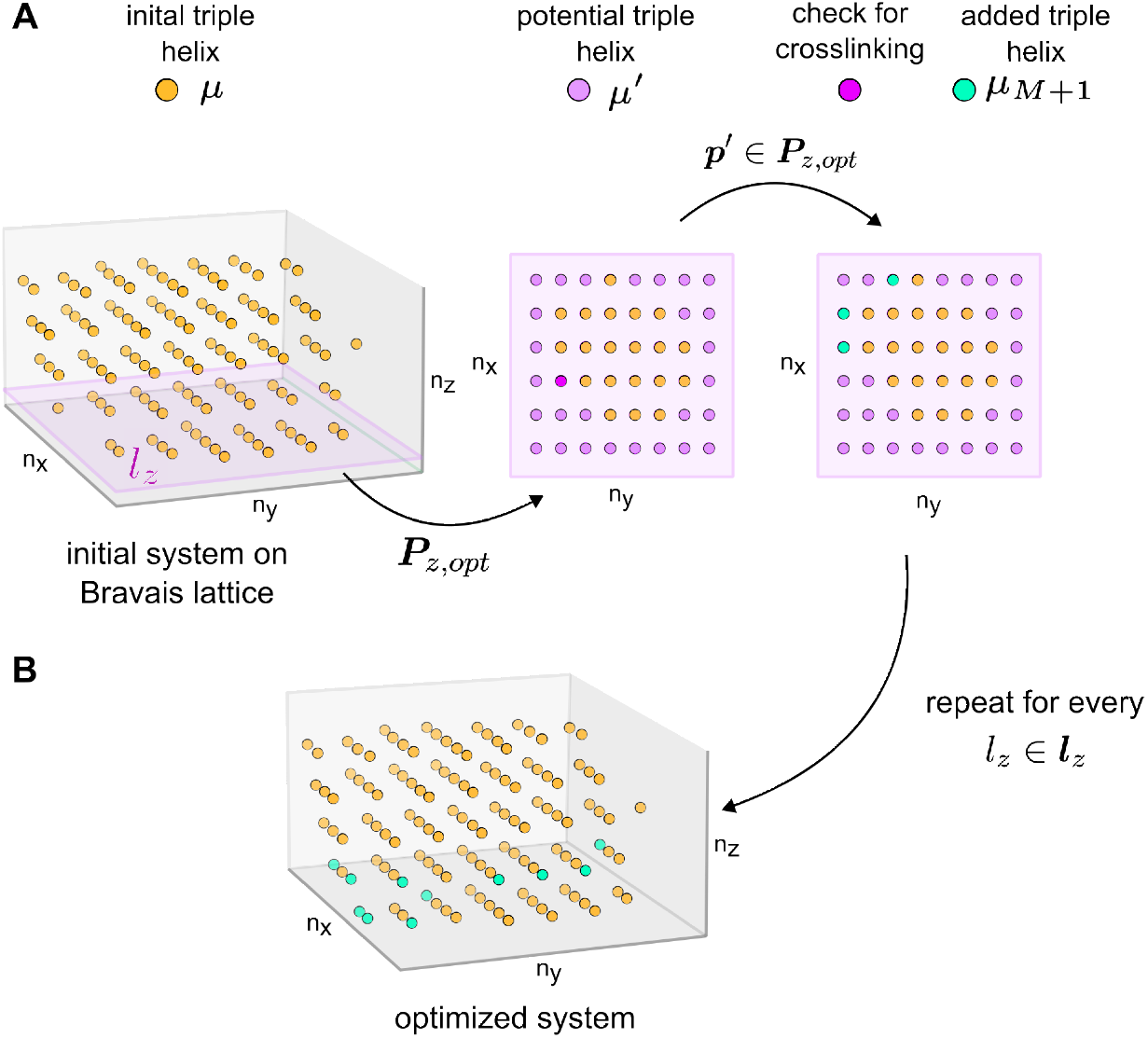
Structural optimization process of a collagen fibril on a Bravais lattice. Starting with an initial system of triple helices *µ* (green points) arranged on a Bravais lattice (A), the process selects a layer *l*_*z*_ for optimization. This layer spans an optimization space ***P***_***m***_ containing potential new triple helices *µ*^*′*^ (light purple). Each potential triple helix *µ*^*′*^ is evaluated for possible crosslinking with existing helices in the model. Triple helices that can be crosslinked (yellow points) are then added to the original system, resulting in an optimized structure (B). This procedure can be repeated for any number of *n*_*x*_, *n*_*y*_−layers, gradually enhancing the fibril’s structural integrity while maintaining its lattice arrangement.

The algorithm proceeds by randomly selecting points ***p***^*′*^∈ **P**_**z**,opt_ and mapping them to Cartesian space to obtain transformation matrices ***T*** ^***′***^ and translation vectors ***t***^*′*^. We translate the model *µ*_1_ crosslink positions by ***t***^*′*^ and check intercrosslink distances between the new potential model *µ*^*′*^ and existing models *µ* (Figure 2A, light purple squares). Contact criteria are defined based on a 0.3 nm cutoff for interatomic distances, which is typical for non-covalent interactions in proteins. If these criteria are met between *µ*^*′*^ and existing models, we incorporate *µ*^*′*^ into the system as *µ*_***M***+1_; otherwise, we discard it. The solution space is updated after each iteration, and the process continues until the current layer’s solution space is exhausted. We then move to the next layer and repeat the process until all layers are optimized (Figure 2B).

The optimized models and transformation matrices are combined using UCSF Chimera’s *matrixset* command to generate an atomistic structure of the collagen microfibril. Post-processing involves cutting each triple helix to the user-specified length (up to 335 nm) and capping termini with neutral Acetyl (ACE) or N-Methyl (NME) groups, preparing the structure for subsequent molecular dynamics simulations.

Our optimization approach resulted in substantial enhancements to the collagen microfibril’s structural properties. We observed a 28% improvement in packing uniformity, quantified by the reduction in point density standard deviation across the lattice. The optimized structures showed excellent agreement with experimental data. We measured an average D-band periodicity of 66.95 ±0.01 nm, closely aligning with the experimentally inferred value of 67 nm. To calculate this periodicity, we applied K-means clustering [21, 22] (k=10) to the z-coordinates of cross-linking residues, identifying distinct bands along the fibril axis. The D-band was then computed as the sum of the gap and overlap distances between adjacent clusters, with distances below 38 nm classified as overlaps and those above as gaps. Additionally, we observed an average lateral molecular spacing of 1.44 ± 0.12 nm, which falls within the range of reported experimental values (1.1 nm for completely dry tissue to 1.8 nm for fresh tissue [23]).

This lateral spacing was calculated by projecting the molecule coordinates onto a plane perpendicular to the fibril’s principal axis, followed by a nearest-neighbor analysis in each quadrant around individual molecules to determine local spacings.

The flexibility of our approach allows for the exploration of different packing configurations by modifying the solution space. For example, considering up to the fourth layer in the *n*_*z*_ dimension (*δ*_*z*_ = 4) results in a Bravais lattice with a slightly different geometrical arrangement of points, particularly for layers closer to the lattice origin. This adaptability enables fine-tuning of the optimization process to meet specific structural requirements or investigate various packing configurations.

Notably, our Bravais lattice-based optimization method demonstrated a tenfold increase in computational efficiency for basic operations compared to traditional Cartesian approaches. In conclusion, our procedure provides an efficient and flexible approach to generate realistic collagen microfibril structures. The resulting models not only exhibit improved structural properties but also align closely with experimental observations, making them valuable for further computational studies of collagen mechanics and function.

### Crosslink Specification for Collagen Microfibrils

Collagen microfibrils are stabilized by intermolecular covalent crosslinks derived from lysine and hydroxylysine residues in the telopeptide regions of the triple helix. These include divalent crosslinks such as hydroxylysine-keto-norleucine (HLKNL) and trivalent crosslinks like hydroxylysyl-pyridinoline (PYD). The overall crosslinking density and the ratio of different crosslink types varies across species, tissues types and age, and is further altered by disease [24]. ColBuilder enables precise control over the composition and density of these crosslinks, accommodating various types and combinations (see Figure S4 in the Supplementary Information for a full list of available structures).

The tool offers two specification methods. First, users can mix triple helices with different crosslink types by defining their ratios. The algorithm then randomly combines these helices to generate a microfibril with the desired composition. Second, ColBuilder can randomly remove existing crosslinks, replacing them with lysine residues using Chimera’s *swapaa* command [19], at a user-defined rate.

Figure 3D shows three different crosslink setups generated based on the Bravais lattice: pure trivalent, trivalent with randomly removed crosslinks, and mixed divalent-trivalent. Trivalent PYD crosslinks (shown in green) are located at the transition between gap and overlap regions (3A), linking adjacent triple helices. The cross-section of this microfibril reveals equally spaced crosslinks forming a rounded shape. The second setup demonstrates the capability of ColBuilder to generate models with varying crosslinking density. We reduced the number of PYD crosslinks within the pure trivalent crosslinked microfibril by randomly replacing 30% of the trivalent crosslinks with lysine residues. This resulted in a microfibril with 70% pure PYD crosslinks, featuring a significantly lower crosslink density compared to the other two microfibrils. For the mixed divalent-trivalent crosslinked microfibril, we defined four types of collagen molecules based on their crosslinks at each telopeptide region: divalent-divalent, trivalent-divalent, divalent-trivalent, and trivalent-trivalent. These were combined in equal proportions, resulting in a microfibril containing both divalent HLKNL (yellow) and trivalent PYD (orange) crosslinks.

**Figure 3.**
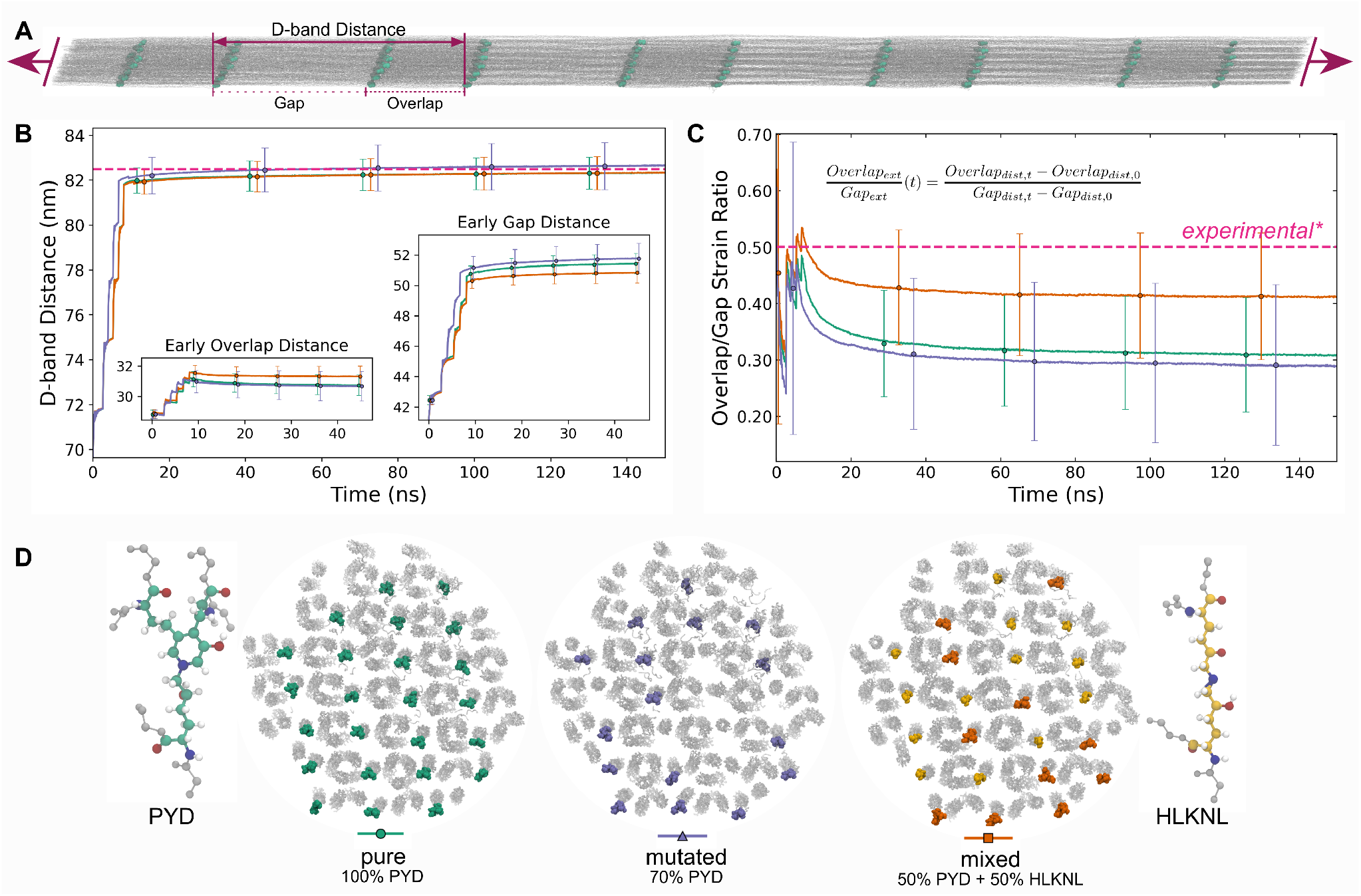
Validation of collagen models with molecular dynamics simulations. (A) Simulation snapshot of a 335 nm-long pulled collagen fibril, illustrating multiple gap and overlap regions with crosslink atoms highlighted as green spheres. (B) D-band distance in nm and (C) overlap/gap strain ratio evolution with simulation time (n sec) for different collagen models illustrated in (D) (insets in (B) show overlap and gap distances early in the simulation, when the fibril was going through greater structural change due to pulling). (D) Cross-sections of collagen fibrils with pure (100% PYD, green), mutated (70% PYD, purple), and mixed (50% PYD + 50% HLKNL, orange) crosslinks. In both B and C, experimentally reported values [39] are shown by the dashed pink line. Error bars represent the standard error of the mean, combining frame-wise variability and trajectory uncertainty, and are horizontally offset for clarity. Error magnitude remains approximately constant throughout the simulation, and line plots show true data. Detailed methods for error calculations are provided in the SI.

These approaches allow for fine-tuning of crosslink density and composition, which are expected to significantly affect the microfibril’s mechanical properties at both macro and microscopic scales [25, 26]. Additionally, users can add their own crosslink structures to ColBuilder’s database, enabling them to model and study the impact of crosslinking variations on fibril structure and mechanics, critical for understanding conditions such as aging and diseases where crosslink patterns and types are altered [27, 28].

### All-Atom Simulations validate models from ColBuilder

To validate the stability and force-extension behavior of the collagen microfibrils generated by ColBuilder, we performed all-atom molecular dynamics simulations. To demonstrate the capabilities of ColBuilder, we chose a fibril length of 335 nm which covers the length of a full triple helix, resulting in 5 complete gap and 4 complete overlap regions. This goes far beyond previous all-atom MD simulations of one gap and one overlap region, the only system size available in our previous database [15]. We generated the microfibril topology using GROMACS 2023 [29], combining the Amber99sb*-ildnp force field [30, 31] for the protein and TIP3P for water with GROMACS’ *pdb2gmx* tool. The simulations of our system of approximately 43 million atoms were conducted using the GENESIS MD engine on the Fugaku supercomputer [32, 33].

Our protocol began with energy minimization in vacuum, followed by solvation and further minimization. We then equilibrated the system through a series of steps: (1) NVT simulation at 300 K over 10 ns using the stochastic velocity rescaling (SVR) [34] with gradually increasing time steps (0.5 to 2 fs), and (2) 5 ns NPT simulation at 1 atm and 300 K using the SRV thermostat and the Martyna-Tobias-Klein (MTK) barostat [35], with gradually increasing time steps (0.5 to 2 fs). Throughout equilibration, backbone atoms were position-restrained (force constant is decreased gradually from 2.5 kcalmol^−1^ Å^−2^ to 1.0 kcalmol^−1^ Å^−2^ in the NVT equilibration and from 1.0 kcalmol^−1^ Å^−2^ to 0.3 kcalmol^−1^ Å^−2^ in the NPT simulation) to maintain microfibril structure while allowing side-chain relaxation. Periodic boundary conditions were applied in all directions. We used group-based thermostat/barostat with optimal temperature evaluation [36, 37].

To allow the 300 nm system to structurally adopt to the applied external force, we implemented a multi-step constant force protocol, incrementally increasing force to 1 nN per strand over 16.5 ns, followed by a 495 ns sustained load simulation. This force magnitude was chosen based on estimated physiological loads on individual collagen molecules, corresponding to stresses in the tens of MP range [38], and the duration was determined to be sufficient for observing initial structural reorganization based on previous studies [9].

Figure 3A shows a snapshot of the collagen microfibril with trivalent PYD crosslinks under the applied force. Notably, no collagen molecule was pulled out during the simulation, demonstrating the structural integrity of our model. The snapshot also reveals differences in molecular packing and stretching between gap and overlap regions under force, with the overlap region showing more parallel alignment of collagen triple helices and less elongation compared to the more intertwined gap region.

We analyzed the mechanical response by measuring three key parameters: end-to-end distance of the entire microfibril, D-band distance - defined as the sum of overlap and gap region lengths -, and the overlap/gap strain ratio - calculated as the ratio between the extensions of overlap and gap regions. These measurements were conducted for three distinct crosslink configurations: pure trivalent, partly trivalent (70% trivalent, 30% absent), and mixed divalent-trivalent crosslinked microfibrils. This approach allowed comparison of their mechanical behaviors under load. To ensure reproducibility and assess variability, we performed three independent simulations for each crosslink configuration.

Our results show that all distance measures increased sharply at the onset of force application before asymptotically approaching constant values. The 335 nm-long microfibril extended by approximately 23%, with the partly trivalent crosslinked microfibril stretching the most (*>*409 nm), followed closely by the pure trivalent and mixed crosslinked microfibrils (≈408 nm) (Figure S3, in the Supplementary Information).

The D-band stretching, resulting from the sum of gap and overlap region elongations, showed similar trends (Figure 3B). We observed elongations of 82.4 nm and 82.8 nm for the pure trivalent/mixed divalent-trivalent and the partly trivalent crosslinked microfibrils, respectively. Notably, this 22% applied strain D-band elongation aligns well with both smaller-scale AA simulations under force (82.5 ± 1.0 nm) and atomic force microscopy (AFM) nanoindentation experiments from the literature (80 to 82.5 nm) [26, 39–41].

The overlap region exhibited a unique stretching pattern (inset of Figure 3B), with an initial sudden increase from 27.9 nm to 31.3 nm, followed by a gradual decrease to constant values of 30.3 nm and 30.7 nm for the mixed divalent-trivalent and the pure/partly trivalent crosslinked microfibrils, respectively. Concurrently, the gap region (also shown in the inset of Figure 3B) expanded from 39.7 nm to 52.1 nm for the partly trivalent and mixed crosslinked microfibrils, and to 51.7 nm for the pure trivalent microfibril. These measurements yielded an average overlap-gap strain ratio of 25% to 30% (Figure 3C), suggesting that the overlap region is approximately three to four times stiffer than the gap region. This finding agrees well with AFM nanoindentation experiments reporting a 25% to 100% higher stiffness in the overlap region compared to the gap region [39, 41].

It is important to note that these mechanical properties can vary depending on factors such as collagen type and tissue of origin (e.g., Achilles or rat tail tendon), experimental conditions (fibril humidity, temperature), and applied force or strain. Despite these potential variations, our collagen microfibrillar structure largely reproduces key structural and mechanical observables reported in the literature, particularly the D-band lengthening and the overlap-gap strain ratio. These results validate the structural integrity of the collagen microfibrils generated by ColBuilder, demonstrating their suitability for further studies of collagen mechanics and function under physiologically relevant conditions.

### Computational Performance

To assess the efficiency of ColBuilder, we analyzed its computational performance in relation to the size and complexity of the generated collagen microfibrils. We measured the wall clock time - the actual time taken to complete a run - for various configurations, focusing on how it scales with the number of collagen chains in the microfibril.

Our analysis revealed a strong correlation between the number of chains in the microfibril and the computational time required. This number of chains is directly influenced by both the contact distance parameter and the desired fibril length, making it a useful metric for predicting performance across different configurations.

Figure S5 illustrates the relationship between the number of collagen chains the wall clock time and the contact distance (dc). We observed that the computational time increases exponentially with the number of chains/contact distance. To provide context for these numbers, we can relate them back to the physical parameters of the microfibril. A typical configuration with a radius of 12 nm (*dc* = 45 Å) and a fibril length of 330 nm resulted in 271 chains and took around 6 minutes 34 seconds to generate. Increasing the contact distance to the double value (*dc* = 90 Å), corresponding to a radius of 16.5 nm while maintaining the same length increased the chain count to 739 and the computational time to 51 minutes and 25 seconds.

These performance metrics were obtained using a standard desktop computer with an Intel Core i5-9600K processor, 3x8 GB of DIMM DDR4 RAM, and an NVIDIA Corporation TU106 GPU, running Ubuntu 22.04.4 LTS. Overall, this desktop computer cannot handle contact distance exceeding 100 Å or radi larger than 17 nm due to memory issues. It is worth noting that runtime may vary depending on the specific hardware configuration used. The observed computational efficiency of ColBuilder makes it a practical tool for researchers, enabling the efficient generation and iteration of collagen microfibril models for various studies in biomechanics and structural biology.

## ColBuilder web server

In addition to the command line application, we provide a web server application of ColBuilder. The web server is implemented within the Flask framework and offers the full functionality of the command line version for selected species and crosslink combinations. ColBuilder web server is freely available at: colbuilder.mpip-mainz.mpg.de.

## Conclusion

We have presented ColBuilder, a computational tool that automates the generation of collagen fibril structures with unprecedented control over their composition and architecture. The integration of homology modeling, higher-order structure generation, and Bravais lattice optimization allows researchers to systematically explore sequence variations, crosslinking patterns, and dimensional parameters that were previously challenging to investigate. The tool’s web interface and command-line application make these capabilities accessible to both structural biologists and biophysics researchers, while its modular design facilitates future extensions.

The ColBuilder project is actively developing. Current work focuses on expanding the crosslink library to include advanced glycation end-product (AGE) crosslinks and implementing topology generation for coarse-grained (Martini) force fields. The demonstrated capability to generate and simulate full-length fibrils establishes a foundation for investigating different collagen types and disease-specific modifications. The current starting point of ColBuilder is a model from X-ray scattering [3]. ColBuilder will greatly profit from additional knowledge on collagen’s hierarchical structure from other sources, being it electron microscopy, new data on crosslinking or other post-translational modifications from mass spectrometry, as well as better insights into the telopeptide packing from AlphaFold [42] or other structure prediction methods. These developments will enable faithful computational studies of collagen mechanics across scales, from molecular interactions to tissue-level properties.

## Author contributions statement

D.M. and M.B. designed and implemented ColBuilder, developed the computational framework, and performed the analysis. J.B. contributed to the software development and testing. V.V., T.S., and M.S. designed and implemented the web server interface. J.J. and Y.S. implemented and performed the molecular dynamics simulations on Fugaku. D.M., M.B., and F.G. wrote the manuscript with input from all authors. F.G. conceived and supervised the project.

## Data Availability Statement

FASTA files for collagen type I of all species discussed in this study are available in the ColBuilder GitHub repository. Code used for computational analyses will be made available in a public GitHub repository upon publication. Simulation data utilized in validation of our models will be made available upon request.

## Acknowledgments

This work was supported by the Klaus Tschira Foundation and the European Research Council (ERC) [grant number 101002812]. D.M. acknowledges funding from the Marie Sklodowska-Curie Actions Individual Fellowship [grant number 101151862]. F.G. acknowledges funding from the Deutsche Forschungsgemeinschaft (DFG, German Research Foundation) under Germany’s Excellence Strategy for the Excellence Cluster “3D Matter Made to Order” (EXC-2082/1-390761711). The authors acknowledge support by the state of Baden-Württemberg through bwHPC for computational resources on the bwForCluster Helix. Additional computational resources were provided by the RIKEN Center for Computational Science on Fugaku (Project ID: ra000003 and Project ID: hp230212). The web server was developed with the help of ChatGPT.

## List of Supplementary Materials

Supplementary text Figures S1 to S5 Tables S1 and S2

## References

1. Karl E Kadler, David F Holmes, John A Trotter, and John A Chapman. Collagen fibril formation. Biochemical Journal, 316(1):1–11, 1996.

2. Matthew D Shoulders and Ronald T Raines. Collagen structure and stability. Annual review of biochemistry, 78(1):929–958, 2009.

3. Joseph PRO Orgel, Thomas C Irving, Andrew Miller, and Tim J Wess. Microfibrillar structure of type i collagen in situ. Proceedings of the National Academy of Sciences, 103(24):9001–9005, 2006.

4. Lynn W Jelinski, CE Sullivan, and DA Torchia. 2h nmr study of molecular motion in collagen fibrils. Nature, 284(5756):531–534, 1980.

5. Paulo De Sa Peixoto, Guillaume Laurent, Thierry Azäis, and Gervaise Mosser. Solid-state nmr study reveals collagen i structural modifications of amino acid side chains upon fibrillogenesis. Journal of Biological Chemistry, 288(11):7528–7535, 2013.

6. Ian Streeter and Nora H de Leeuw. Atomistic modeling of collagen proteins in their fibrillar environment. The Journal of Physical Chemistry B, 114(41):13263–13270, 2010.

7. Susanna Monti, Simona Bronco, and Chiara Cappelli. Toward the supramolecular structure of collagen: a molecular dynamics approach. The Journal of Physical Chemistry B, 109(22):11389–11398, 2005.

8. Baptiste Depalle, Zhao Qin, Sandra J. Shefelbine, and Markus J. Buehler. Influence of cross-link structure, density and mechanical properties in the mesoscale deformation mechanisms of collagen fibrils. Journal of the Mechanical Behavior of Biomedical Materials, 52:1–13, 2015. SI:Collagen mechanics.

9. Christopher Zapp, Agnieszka Obarska-Kosinska, Benedikt Rennekamp, Markus Kurth, David M Hudson, Davide Mercadante, Uladzimir Barayeu, Tobias P Dick, Vasyl Denysenkov, Thomas Prisner, et al. Mechanoradicals in tensed tendon collagen as a source of oxidative stress. Nature communications, 11(1):2315, 2020.

10. Shiamalee Perumal, Olga Antipova, and Joseph PRO Orgel. Collagen fibril architecture, domain organization, and triple-helical conformation govern its proteolysis. Proceedings of the National Academy of Sciences, 105(8):2824–2829, 2008.

11. Ian Streeter and Nora H. de Leeuw. A molecular dynamics study of the interprotein interactions in collagen fibrils. Soft Matter, 7:3373–3382, 2011.

12. Jumin Lee, Dhilon S Patel, Jonas Ståhle, Sang-Jun Park, Nathan R Kern, Seonghoon Kim, Joonseong Lee, Xi Cheng, Miguel A Valvano, Otto Holst, et al. Charmm-gui membrane builder for complex biological membrane simulations with glycolipids and lipoglycans. Journal of chemical theory and computation, 15(1):775–786, 2018.

13. Leandro Martínez, Ricardo Andrade, Ernesto G Birgin, and José Mario Martínez. Packmol: A package for building initial configurations for molecular dynamics simulations. Journal of computational chemistry, 30(13):2157–2164, 2009.

14. Yifei Qi, Jumin Lee, Jeffery B Klauda, and Wonpil Im. Charmm-gui nanodisc builder for modeling and simulation of various nanodisc systems. Journal of computational chemistry, 40(7):893–899, 2019.

15. Agnieszka Obarska-Kosinska, Benedikt Rennekamp, Aysecan Ünal, and Frauke Gräter. Colbuilder: A server to build collagen fibril models. Biophysical Journal, 120(17):3544–3549, 2021.

16. Peter J. A. Cock, Tiago Antao, Jeffrey T. Chang, Brad A. Chapman, Cymon J. Cox, Andrew Dalke, Iddo Friedberg, Thomas Hamelryck, Frank Kauff, Bartek Wilczynski, and Michiel J. L. de Hoon. Biopython: freely available Python tools for computational molecular biology and bioinformatics. Bioinformatics, 25(11):1422–1423, 03 2009.

17. Robert C. Edgar. MUSCLE: multiple sequence alignment with high accuracy and high throughput. Nucleic Acids Research, 32(5):1792–1797, 03 2004.

18. Jordi Bella, Mark Eaton, Barbara Brodsky, and Helen M Berman. Crystal and molecular structure of a collagen-like peptide at 1.9 å resolution. Science, 266(5182):75–81, 1994.

19. Eric F. Pettersen, Thomas D. Goddard, Conrad C. Huang, Gregory S. Couch, Daniel M. Greenblatt, Elaine C. Meng, and Thomas E. Ferrin. UCSF Chimera—A visualization system for exploratory research and analysis. 25(13):1605–1612.

20. Dennis W. Bennett. Understanding Single-Crystal X-ray Crystallography. Wiley-VCH-Verl, 2010.

21. J Macqueen. Some methods for classification and analysis of multivariate observations. In Proceedings of 5-th Berkeley Symposium on Mathematical Statistics and Probability/University of California Press, 1967.

22. Fabian Pedregosa, Gäel Varoquaux, Alexandre Gramfort, Vincent Michel, Bertrand Thirion, Olivier Grisel, Mathieu Blondel, Peter Prettenhofer, Ron Weiss, Vincent Dubourg, et al. Scikit-learn: Machine learning in python. the Journal of machine Learning research, 12:2825–2830, 2011.

23. P Fratzl, N Fratzl-Zelman, and K Klaushofer. Collagen packing and mineralization. an x-ray scattering investigation of turkey leg tendon. Biophysical journal, 64(1):260–266, 1993.

24. Gautieri Alfonso Snedeker, Jess G. The role of collagen crosslinks in ageing and diabetes - the good, the bad, and the ugly. Muscles Ligaments Tendons J, 4:303–8, 2014.

25. Albert L Kwansa, Raffaella De Vita, and Joseph W Freeman. Tensile mechanical properties of collagen type i and its enzymatic crosslinks. Biophysical chemistry, 214:1–10, 2016.

26. Benedikt Rennekamp, Christoph Karfusehr, Markus Kurth, Aysecan Ünal, Debora Monego, Kai Riedmiller, Ganna Gryn’ova, David M Hudson, and Frauke Gräter. Collagen breaks at weak sacrificial bonds taming its mechanoradicals. Nature Communications, 14(1):2075, 2023.

27. Vincent M Monnier, Marcus Glomb, Abdelhamid Elgawish, and David R Sell. The mechanism of collagen cross-linking in diabetes: a puzzle nearing resolution. Diabetes, 45(Supplement 3):S67–S72, 1996.

28. Jess G Snedeker and Alfonso Gautieri. The role of collagen crosslinks in ageing and diabetes-the good, the bad, and the ugly. Muscles, ligaments and tendons journal, 4(3):303, 2014.

29. David Van Der Spoel, Erik Lindahl, Berk Hess, Gerrit Groenhof, Alan E Mark, and Herman JC Berendsen. Gromacs: fast, flexible, and free. Journal of computational chemistry, 26(16):1701–1718, 2005.

30. Robert B Best and Gerhard Hummer. Optimized molecular dynamics force fields applied to the helix- coil transition of polypeptides. The journal of physical chemistry B, 113(26):9004–9015, 2009.

31. Kresten Lindorff-Larsen, Stefano Piana, Kim Palmo, Paul Maragakis, John L Klepeis, Ron O Dror, and David E Shaw. Improved side-chain torsion potentials for the amber ff99sb protein force field. Proteins: Structure, Function, and Bioinformatics, 78(8):1950–1958, 2010.

32. Jaewoon Jung, Kiyoshi Yagi, Cheng Tan, Hiraku Oshima, Takaharu Mori, Isseki Yu, Yasuhiro Matsunaga, Chigusa Kobayashi, Shingo Ito, Diego Ugarte La Torre, et al. Genesis 2.1: High-performance molecular dynamics software for enhanced sampling and free-energy calculations for atomistic, coarse-grained, and quantum mechanics/molecular mechanics models. The Journal of Physical Chemistry B, 2024.

33. Jaewoon Jung, Chigusa Kobayashi, Kento Kasahara, Cheng Tan, Akiyoshi Kuroda, Kazuo Minami, Shigeru Ishiduki, Tatsuo Nishiki, Hikaru Inoue, Yutaka Ishikawa, et al. New parallel computing algorithm of molecular dynamics for extremely huge scale biological systems. Journal of computational chemistry, 42(4):231–241, 2021.

34. Giovanni Bussi and Michele Parrinello. Accurate sampling using langevin dynamics. Physical review E-Statistical, Nonlinear, and Soft Matter Physics, 75:056707, 2007.

35. Glenn J Martyna, Douglas J Tobias, and Michael L Klein. Constant pressure molecular dynamics algorithms. The Journal of chemical physics, 101(5):4177–4189, 1994.

36. Jaewoon Jung, Chigusa Kobayashi, and Yuji Sugita. Optimal temperature evaluation in molecular dynamics simulations with a large time step. Journal of Chemical Theory and Computation, 15(1):84–94, 2018.

37. Jaewoon Jung and Yuji Sugita. Group-based evaluation of temperature and pressure for molecular dynamics simulation with a large time step. The Journal of Chemical Physics, 153(23), 2020.

38. Paavo V Komi. Relevance of in vivo force measurements to human biomechanics. Journal of biomechanics, 23:23–34, 1990.

39. Chris J Peacock and Laurent Kreplak. Nanomechanical mapping of single collagen fibrils under tension. Nanoscale, 11(30):14417–14425, 2019.

40. Emilie Gachon and Patrick Mesquida. Stretching single collagen fibrils reveals nonlinear mechanical behavior. Biophysical Journal, 118(6):1401–1408, 2020.

41. Majid Minary-Jolandan and Min-Feng Yu. Nanomechanical heterogeneity in the gap and overlap regions of type i collagen fibrils with implications for bone heterogeneity. Biomacromolecules, 10(9):2565–2570, 2009.

42. Evans Richard Pritzel Alexander Green Tim Figurnov Michael Ronneberger Olaf Tunyasuvunakool Kathryn Bates Russ Žídek Augustin Potapenko Anna Bridgland Alex Meyer Clemens Kohl Simon A.A. Ballard Andrew J. Cowie Andrew Romera-Paredes Bernardino Nikolov Stanislav Jain Rishub Adler Jonas Back Trevor Petersen Stig Reiman David Clancy Ellen Zielinski Michal Steinegger Martin Pacholska Michalina Berghammer Tamas Bodenstein Sebastian Silver David Vinyals Oriol Senior Andrew W. Kavukcuoglu Koray Kohli Pushmeet Hassabis Demis Jumper, John. Highly accurate protein structure prediction with alphafold. Nature, 596(7873):583––589, 2021.

