## Supplementary Information for "ColBuilder: Flexible structure generation of crosslinked collagen fibrils"

#### Homology Modeling and Initial Structure Generation

Collagen triple helices for various species were modeled using ColBuilder, with *Rattus novergicus* collagen type I as the reference. Sequence alignment was performed to assess conservation, and per-residue RMSD values were calculated to quantify structural differences. As shown in Figure S1, the RMSD values for the human and rat collagen chains remain below 1.5 Å, indicating a high level of structural accuracy in our homology models.

To further evaluate the consistency of the models across species, Table S1 shows the average sequence identity, per-residue RMSD, and axial rise per triplet for collagen triple helices generated for all species in our database. These values demonstrate consistent homology model quality, with RMSD values typically ranging between 1.0-1.4 Å, supporting the reliability of our homology modeling procedure across a diverse set of species.

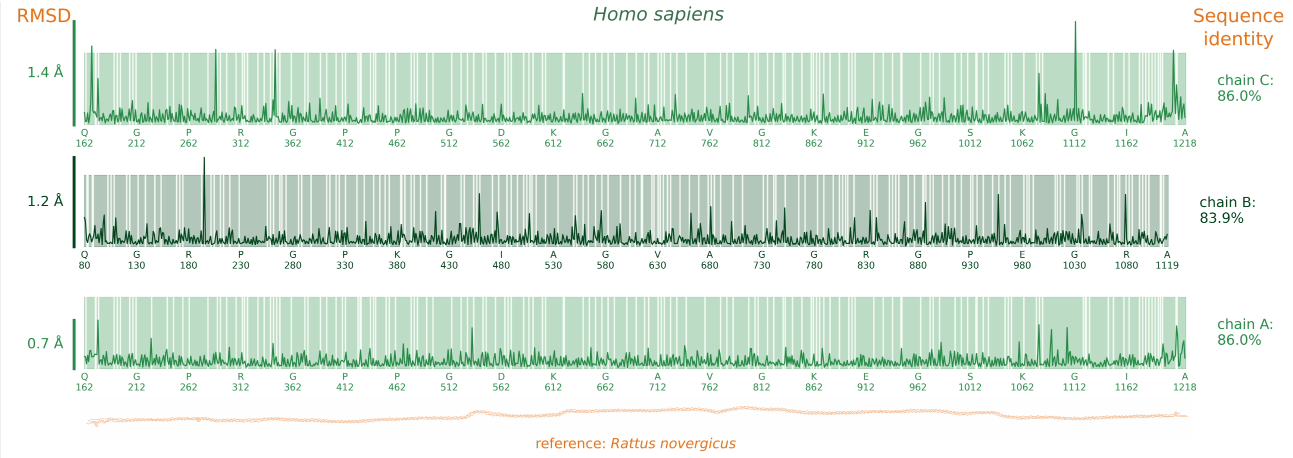


**Figure S1.** Sequence alignment (solid color: identical, and gaps: diverging residues) and per-residue RMSD (line) between each chain of human (*Homo sapiens*) and rat (*Rattus novergicus*) collagen type I, shown at the bottom for reference. The magnitude of the RMSD (Å) is shown on the left, and the average sequence identity for each chain on the right.

**Table S1.** Average sequence identity, per-residue RMSD ± SD, and average axial rise per triplet ± SD for collagen triple helices generated with ColBuilder. Collagen type I of *Rattus novergicus* was used as a reference.

| Species | Sequence Identity, % | RMSD, Å | Axial Rise per Triplet, Å |
|---|---|---|---|
| *Otolemur garnettii* | 93.57 | 1.25 ± 0.32 | 9.29 ± 0.46 |
| *Xiphophorus maculatus* | 72.01 | 1.39 ± 0.10 | 9.26 ± 0.45 |
| *Tetraodonni groviridis* | 71.6 | 1.35 ± 0.09 | 9.24 ± 0.37 |
| *Oreochromis niloticus* | 75.04 | 1.29 ± 0.08 | 9.28 ± 0.45 |
| *Callithrix jacchus* | 94.81 | 1.22 ± 0.3 | 9.27 ± 0.44 |
| *Pongo abelii* | 93.55 | 1.25 ± 0.17 | 9.28 ± 0.45 |
| *Pelodiscus sinensis* | 83.29 | 1.24 ± 0.16 | 9.23 ± 0.38 |
| *Pan troglodytes* | 94.72 | 1.08 ± 0.28 | 9.28 ± 0.44 |
| *Mustela putorius* | 95.32 | 1.06 ± 0.29 | 9.27 ± 0.44 |
| *Canis lupus* | 95.29 | 1.08 ± 0.28 | 9.28 ± 0.44 |
| *Ailuropoda melanoleuca* | 93.74 | 1.28 ± 0.13 | 9.27 ± 0.50 |
| *Homo sapiens* | 85.3 | 1.12 ± 0.32 | 9.28 ± 0.44 |
| *Danio rerio* | 75.43 | 1.23 ± 0.01 | 9.28 ± 0.46 |
| *Mus musculus* | 97.64 | 1.12 ± 0.26 | 9.29 ± 0.46 |
| *Loxodonta africana* | 90.17 | 1.22 ± 0.20 | 9.25 ± 0.43 |
| *Bos taurus* | 94.49 | 1.08 ± 0.27 | 9.3 ± 0.44 |
| *Oryzias latipes* | 75.04 | 1.29 ± 0.08 | 9.28 ± 0.45 |
| *Myotis lucifugus* | 94.5 | 1.08 ± 0.28 | 9.29 ± 0.46 |

### Higher-Order Crystal Structural Generation

##### Contact distance parameter

The radius of collagen fibrils was determined using a geometric approach based on the spatial distribution of atoms in the PDB files. The x and y coordinates of all atoms were extracted to create a 2D projection of the fibril structure. A convex hull was then computed from these coordinates using SciPy's *ConvexHull* function, representing the outer boundary of the fibril cross-section. The centroid of all atomic coordinates was calculated as the mean of all x and y coordinates. Distances from this centroid to each point on the convex hull were computed, and the radius of the fibril was defined as the 95th percentile of these distances. This approach was chosen to provide a robust estimate of the fibril's outer boundary while accounting for potential outliers. To quantify the uncertainty in the radius calculation, we used the interquartile range (IQR) of the centroid-to-hull distances. The radius error was calculated as half of the IQR, offering a conservative estimate of the measurement uncertainty that is resistant to outliers. This method allows for a robust estimation of fibril radius that is less sensitive to local irregularities in the atomic coordinates, while providing a reliable measure of the variability in the fibril's cross-sectional shape.

##### All-atom simulations

The end-to-end distance was calculated as the Euclidean distance between the mean positions of the ACE (N-terminal) and NME (C-terminal) cap residues. This measure represents the overall extension of the collagen molecule.

The error bars in Figure 3 B and C in the main text represent the standard error of the mean for D-band distances and overlap/gap strain ratios in our collagen molecular dynamics simulations. We used K-means clustering to identify cross-linking residues and determine overlap and gap regions for each simulation frame. Standard errors for overlap and gap measurements were calculated from the spread of residue positions within clusters. For overlap/gap strain ratios, we propagated these errors using standard techniques. D-band distance errors, in turn, combine two components: (1) the propagated frame-wise overlap and gap errors, and (2) a bootstrap error estimate. The latter used a 10-frame sliding window with 1000 resamples per window. We combined these components using a root-mean-square method. This approach captures both local configurational variability and broader trajectory fluctuations, providing an integrated representation of uncertainty in our simulated collagen systems.

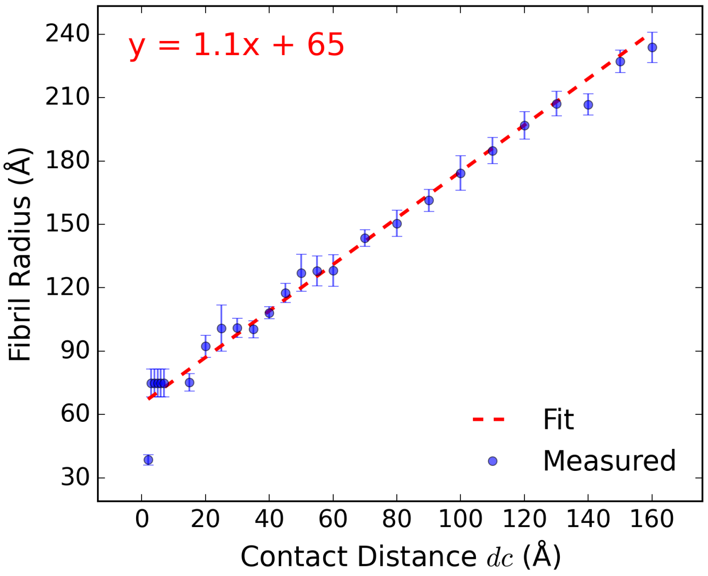


**Figure S2.** Dependence of fibril radius (Å) with ColBuilder.0 parameter contact distance $dc$ (Å). Users can determine the approximate $dc$ value needed to produce a fibril of a given radius by solving for x in the regression equation for the fitting.

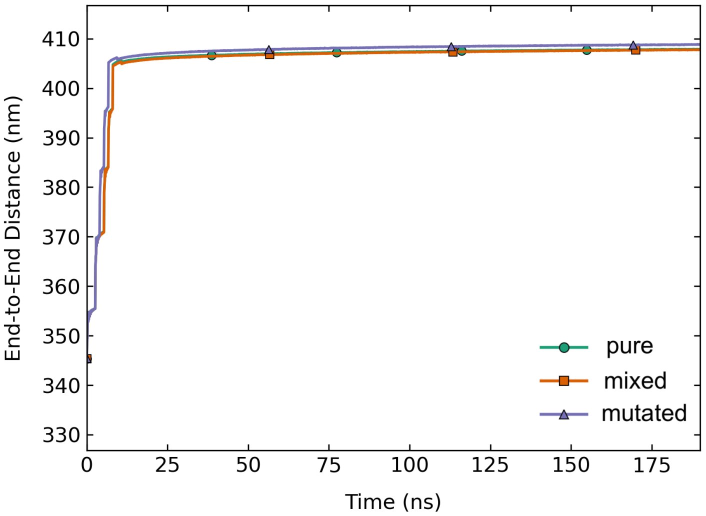


**Figure S3.** Overall extension of collagen fibril, represented by the end-to-end distance (nm) for pure (green, 100% PYD crosslinks), mutated (purple, 70% PYD) and mixed (orange, 50% PYD and 50% HLKNL) collagen fibrils. End-to-end distances were calculated as the Euclidian distance between the mean positions of the ACE (N-terminal) and NME (C-terminal) cap residues.

### Crosslinks

ColBuilder uses a standardized format to define crosslinks between collagen molecules. Each crosslink is specified by a combination of its chemical type (4), the residues involved, and their precise connection points at the atomic level. The definition includes information about the original residue (typically lysine), its terminal location (N or C), the specific residue positions involved in the crosslink, and the detailed atomic connections. For trivalent crosslinks like PYD, three connection points are specified, while divalent crosslinks like HLKNL require only two. Table 2 shows examples of these definitions for *Homo sapiens*, illustrating the format for different crosslink types.

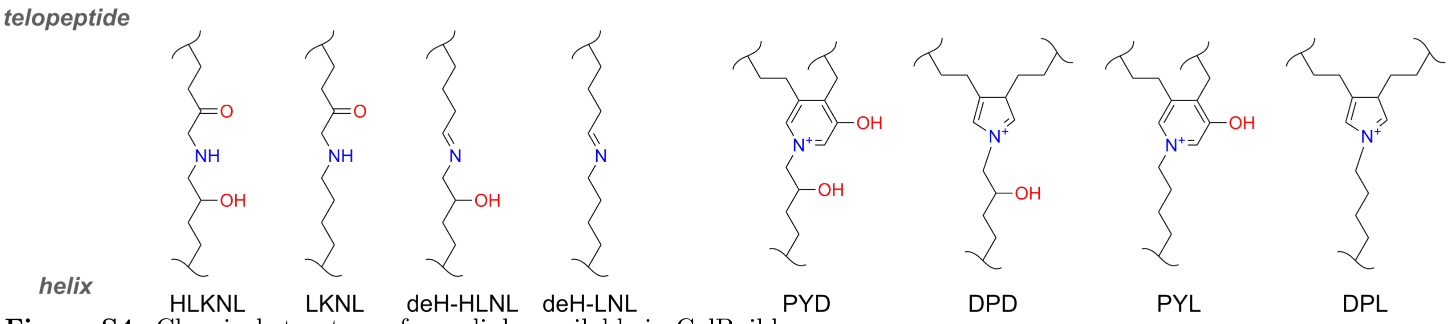


**Figure S4.** Chemical structure of crosslinks available in ColBuilder.

**Table S2.** Example of crosslink definitions in ColBuilder for *Homo sapiens*. Fields are: OR (original residue), terminal (N/C terminal), combination (residue positions involved), type (crosslink type), R1/A1/P1 (first residue/atom/position), R2/A2/P2 (second residue/atom/position), and R3/A31/A32/P3 (third residue/atoms/position for trivalent crosslinks).

| OR | term | combination | type | R1 | A1 | P1 | R2 | A2 | P2 | R3 | A31 | A32 | P3 |
|---|---|---|---|---|---|---|---|---|---|---|---|---|---|
| LYS | C | 1047.C-1047.A-104.C | PYD | LY3 | CG | 1047.C | LY2 | CB | 1047.A | LYX | C13 | C12 | 104.C |
| LYS | N | 9.C-5.B-944.B | PYD | LY3 | CG | 9.C | LY2 | CB | 5.B | LYX | C13 | C12 | 944.B |
| LYS | C | 1047.C-104.C | HLKNL | L5Y | NZ | 1047.C | L4Y | CE | 104.C | NONE | NONE | NONE | NONE |
| LYS | N | 5.B-944.B | HLKNL | L5Y | NZ | 5.B | L4Y | CE | 944.B | NONE | NONE | NONE | NONE |
| LYS | C | 1047.C-104.C | NOCROSS | NONE | NONE | NONE | NONE | NONE | NONE | NONE | NONE | NONE | NONE |

Notes: PYD = pyridinoline (trivalent), HLKNL = hydroxylysinonorleucine (divalent), NOCROSS = no crosslink. Positions are given in residue.chain format. For trivalent crosslinks, R3/A31/A32/P3 specify the third connection point (A1 binds to A31 and A2 binds to A32, while divalent crosslinks use NONE for these fields.

### Computational Performance

We evaluated ColBuilder's computational efficiency by measuring wall clock time as a function of system size, characterized by both contact distance (dc) and the resulting number of triple helices. As shown in Figure 5, the computational time increases exponentially with contact distance, reflecting the growing complexity of structure generation and optimization as more helices are included in the fibril. System Specifications:
Processor: Intel Core i5-9600K processor, CPU family: 6, Model:158, Thread(s) per core: 1, Core(s) per socket: 6, Socket(s): 1, Stepping: 12, @ 3.70GHz
RAM: 3x DIMM DDR4 Synchronous Unbuffered ,size: 8GB, width: 64 bits,clock: 2666MHz (0.4ns)
GPU: NVIDIA Corporation TU106 GPU (NVIDIA-SMI 550.107.02 ), running Ubuntu 22.04.4 LTS

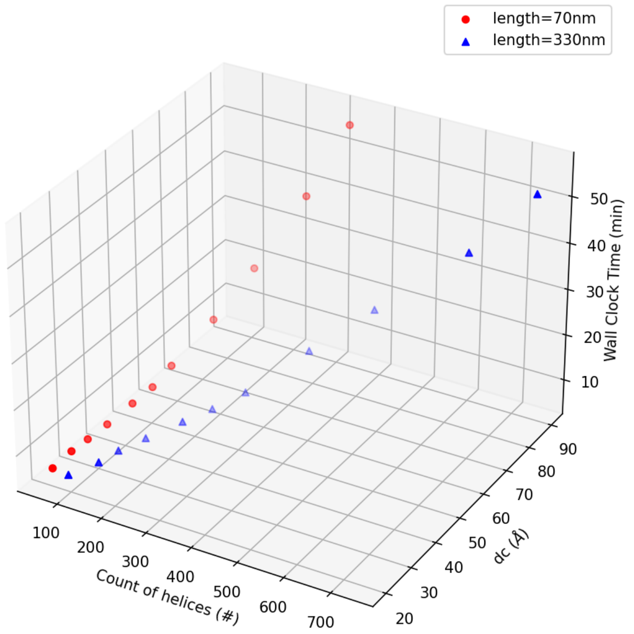


**Figure S5.** The wall clock time vs. the contact distance dc and count of helices. The figure shows the exponential behavior of the time vs. the contact distance for both shown lengths of fibrils.
